# GerAB Residues Predicted to Interact with Water Based on MD Simulations Mediate Germinosome Stability in *Bacillus subtilis* spores

**DOI:** 10.64898/2026.01.10.698761

**Authors:** Longjiao Chen, Silke van Buuren, George Korza, Jocelyne Vreede, Peter Setlow, Stanley Brul

## Abstract

Some species in the *Bacillales* and *Clostridiales* orders form spores under unfavourable environmental conditions. These spores are metabolically dormant and highly resistant to extreme stress. The spore core—analogous to the protoplast of vegetative cells—contains only 25–45% water by wet weight, compared to ~80% in vegetative cells. Upon activation by small-molecule nutrients, spores germinate, restoring their core water content, restoring metabolism and becoming easy to kill, while progressing through outgrowth to vegetative growth. GerAB is the B subunit of the prototypical *Bacillus subtilis* GerA GR (germinant receptor), a membrane protein belonging to the Amino Acid—Polyamine-Organocation (APC) superfamily of transporters. It functions as the L-alanine sensor that initiates germination and was previously predicted, based on molecular dynamics (MD) simulations, to contain a putative water channel. Using MD simulations, we identified low amount of water permeating through GerAB (ranging from 1-121 water molecule/µs in 10 parallel MD simulations), thus revealing a water pathway in GerAB that diverges from the L-alanine binding pocket, suggesting that water transport may play roles in germination beyond facilitating ligand binding. Analysis of water–residue contact frequencies identified eight hydrophilic residues lining this path. Individual substitution of high-contact residues with similarly sized non-polar residues impaired L-alanine germination and disrupted GerAB structural integrity as assessed by Western Blotting. These mutants also respond to the AGFK germinant mixture (L-asparagine, D-glucose, D-fructose and potassium) in slower, yet individually distinct kinetics compared to that of wt spores. These findings prove that water contact residues in GerAB predicted by MD simulations are crucial for the stability of this protein and thus the germinosome complex with all GRs.

## 1 Introduction

Certain species within the *Bacillales* and *Clostridiales* orders can form dormant spores in response to harsh environmental conditions. These spores are metabolically inactive and exhibit exceptional resistance to a wide range of extreme treatments. However, potentially, spores from species such as *Bacillus cereus, Bacillus anthracis*, and *Clostridioides difficile* can cause food spoilage, and foodborne illnesses ^1^. Consequently, the mechanisms underlying spore germination in these bacteria have been investigated over the past decades, especially so since spores lose their extreme resistance properties upon germination. A characteristic feature of bacterial spores is their low core water content. In *Bacillus subtilis*, the spore core—analogous to the cytoplasm of vegetative cells—contains only 25–45% water by wet weight, compared to approximately 80% in vegetative cells. Upon activation by small-molecule nutrients known as germinants, spores initiate germination, during which rapid water uptake occurs. This water influx leads to roughly a two-fold expansion of the spore core volume, a process that takes around 10 minutes in an individual spore ^1–3^. Full hydration of the core marks the completion of germination and enables the resumption of metabolic activity and macromolecular synthesis. Despite its importance, the molecular mechanism by which water is taken up into the spore core during germination remains unknown in spore-forming bacteria. ^1^

During the germination process described above, GRs play a central role by sensing germinants and initiating the molecular cascade that leads to spore revival. There are three functional GRs in the model organism *Bacillus subtilis*, GerA, GerB and GerK, with GerA responding to L-alanine while GerB and GerK respond to a mixture of AGFK (L-asparagine, D-glucose, D-fructose and potassium) in a cooperative manner ^1^. Each GR is composed of three essential subunits, and it has been well-established that all three subunits are indispensable for forming a functional receptor^4^. The three GRs further colocalize with other germination proteins in a protein complex termed a germinosome, in which each component is structurally and functionally dependent on each other^4,5^. Recent studies have significantly advanced our understanding of GR function by showing that the three subunits of GerA assemble into a heteropentameric ion channel, where the GerAA subunit forms a pentameric structure that facilitates the release of monovalent cations upon detection of the germinant L-alanine bound to the GerAB subunit ^6,7^. These findings were made possible by a combination of AlphaFold structure predictions and mutagenesis experiments. Beyond the nutrient sensing function of the B subunit, Blinker *et al*. identified a putative water channel within GerAB using molecular dynamics (MD) simulations, suggesting a novel mechanism for water transport during germination. More recently, Chen *et al*^8^ further characterized this water channel by identifying specific residues in GerAB that form a blockade for water crossing in steered MD simulations. These studies point to a previously less understood role for GerAB in facilitating core hydration, a vital step in spore germination. However, experimental validation, the presumed regulation of this water channel and its precise physiological role in spore germination remain open questions. Given its possible multifunctionality, GerAB makes a fascinating target for elucidating the molecular mechanisms of spore germination.

As no structure has been determined for GerAB with experimental methods, several studies have sought to deepen the understanding of its function with bioinformatics approaches. These investigations have reached a consensus that GerAB is a member of the membrane transporter protein superfamily known as the Five-Helix Inverted Repeat Superfamily. More specifically, GerAB belongs to the Amino Acid-Polyamine-Organocation (APC) superfamily ^7,9,10^. Notably, growing evidence points to the critical role of water in the function of proteins within this superfamily. This holds true for two prominent families within the Five-Helix inverted repeat superfamily, the Amino Acid-Polyamine-Organocation (APC) family and the Neurotransmitter-Sodium Symporter (NSS) family. For example, a well-characterized APC transporter, GkApcT from *Geobacillus kaustophilus*, has been shown to transport water molecules *in silico* with MD simulations. In the absence of quantitative measurements of GkApcT permeability, the study indicates that its water permeability is lower than that of dedicated aquaporins, which exhibit water transport rates on the order of 10^9^s^−1^in AQP1^11^. In GkApcT, water facilitates substrate translocation by acting as a solvent medium, stabilizing nearby residues, and supporting conformational changes required for transport ^12^. Similarly, in the NSS family, the sodium-galactose symporter vSGLT functions not only as a transporter of galactose but also mediates water transport, acting simultaneously as a passive water channel and an active transporter in MD simulations ^13^. Importantly, *B. subtilis* GerAB is thought to share a high degree of structural homology with GkApcT ^7^ and LeuT, another well-studied NSS family transporter^14,15^. Additionally, molecular dynamics simulations have identified a putative water channel within GerAB ^10^, with a pore radius of 0.1-0.3nm throughout 5 parallel 100ns simulations at all times. This supports the hypothesis that water may play a functional role in GerAB. Unlike canonical transporters, GerAB and its homologs may act as *transceptors*—proteins that sense external ligands and trigger downstream signalling without necessarily transporting the ligand itself ^16,17^. Nonetheless, given the critical role water plays in structurally similar transporters, it seemed worthwhile to explore if water transport is also a function of GerAB, thus providing new insight of the role of this GR subunit in spore germination.

To further investigate the mechanism of GerAB’s interaction with water molecules and the role of hydration in spore germination, we performed molecular dynamics simulations of the GerAB–membrane system. These simulations revealed water crossing through GerAB at a low level (ranging from 1 to 121 molecules/µs in 10 parallel MD simulations). Analysing contact frequency between GerAB residues and crossing water molecules identified eight hydrophilic residues that frequently interact with water during crossing events. These residues span the GerAB protein on TM regions 1, 2, 3, 6 and 10, and diverge from the previously revealed L-alanine binding pocket ^7^. To experimentally verify the functional role of these residues, we substituted each one individually with similarly sized hydrophobic amino acid residues and prepared mutant spores. In germination assays, all of the GerAB mutant spores showed defects in L-alanine induced germination. Furthermore, Western Blot revealed impaired GerAB structural integrity. In addition, AGFK induced germination in GerAB mutant spores displayed decreased and individually varied kinetics compared to wt spores. These results indicated that different mutations in GerAB introduce different structural perturbations that eventually affect the function of GerB and GerK GR in various ways. Together, these results demonstrate that the predicted water contact residues in GerAB are structural hotspots in GerA and germinosome stability.

## 2 Result

### 2.1 Low amount of water passage occurred during MD simulations

The RMSD of protein Cα atoms indicated that the structure stabilized around 6 Å after approximately 200 ns, suggesting the system had reached equilibrium beyond this time point (Figure S1A). Cα RMSF analysis across ten independent simulations showed consistent trends: low fluctuations in the transmembrane helical regions and high fluctuations in loop regions (Figure S1B). Based on these results, all analyses were conducted using data from 200 ns onward. One typical water crossing event is captured in Figure 1A as snapshots of MD simulations, and the trajectory for this water molecule is projected on the simulation box in Figure 1B. The overall water permeation profiles from all ten production runs are summarized in Figure 1C. Across the ten simulations, two distinct patterns of water permeation emerged. In nine of the runs, water passage through the channel remained limited (between 1 to 24 molecules/µs), whereas one trajectory exhibited higher amount of water passage at 121 molecules/µs. Moreover, during all simulation trajectories, the permeation events happen in both directions, meaning water could enter from both sides of the protein. With the Z axis of the simulation box aligning with the principal axis of the protein, water entering from low Z coordinate and exiting from high Z coordinate was defined as “forward”, while the other direction was defined as “reverse” (Figure 1C). Throughout the simulations, the number of forward and reverse permeation events are at the same order of magnitude at all times (Figure 1D). This observation is expected, as the simulations were performed under equilibrium conditions with no concentration gradient across the membrane. Consequently, there is no driving force for water entering the protein other than diffusion. In other words, during the simulations, water movement in the “forward” direction is equally likely as movement in the “reverse” direction. The distribution of the time individual water molecules took to pass through GerAB peaks at 5ns and exhibits a long tail extending to 20ns (Figure S2).

**Figure 1.**
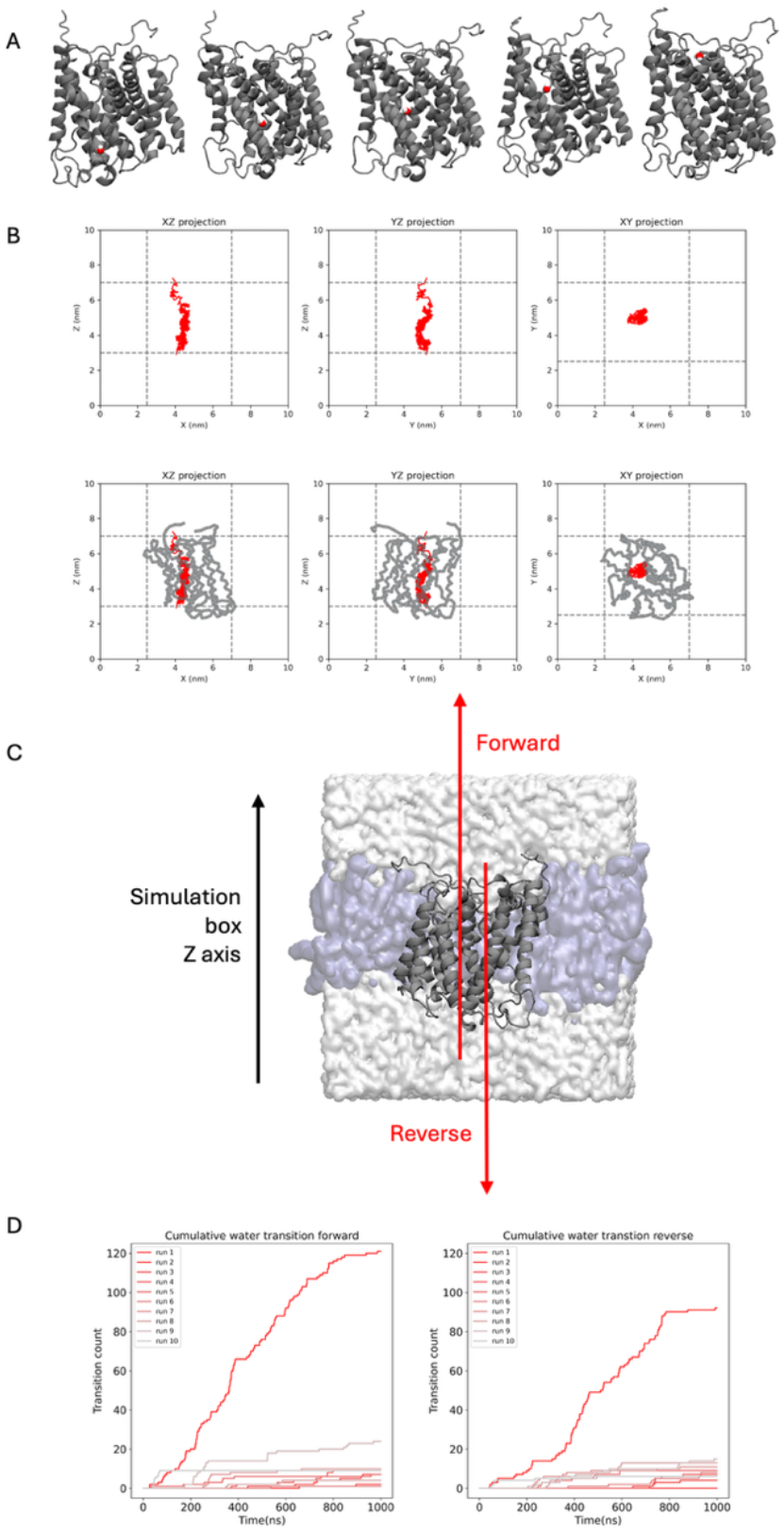
Water passage through GerAB during MD simulations. (A) One typical water passage event shown as snapshots during MD simulations. GerAB protein is coloured in grey and illustrated in New Cartoon. The water molecule passing GerAB was illustrated in space filling, with the Oxygen atom in red and hydrogen atoms in white. (B) Projection of the same water molecule passing through GerAB trajectory on XZ, YZ and XY planes of the simulation box. Three projections of the water trajectories were shown to demonstrate that the water passages selected by our analysis indeed occur through GerAB, rather than penetrating the membrane. The top panel shows the water trajectory. The lower panel shows the water trajectory with protein backbone illustrated in grey thread. Dotted lines indicated the X, Y and Z bound protein to calculate water passage. (C) Simulation box of the protein/membrane system in this study. Water and ions are depicted as a transparent surface, and the lipid bilayer is depicted as blue surface. The direction of the Z axis is shown as a black arrow, and the forward and reverse water passage directions are shown as red arrows. (D) Cumulative water passage profile of ten individual simulation runs in both directions.

### 2.2 Identification of residues interacting with crossing water

To investigate the mechanism of water passing through GerAB, we identified residues that came into contact with water molecules during passing events that occurred in the simulations. The contact frequency *f* of each residue was computed by counting the number of frames where the passing water oxygen was within 3.5 Å (length of a typical hydrogen bond) of any heavy atom of the residue and divided by the total number of frames. We identified 16 residues that exhibited *f* above the statistical significance threshold of 0.01 in both passage directions (Table S1). In the forward water passage direction, all passage water took in a total of 1,285,813 simulation frames to traverse GerAB. A contact frequency of 0.01 of one residue therefore corresponds to 12,858 frames, indicating that the residue was in contact with water for approximately 25.7 ns in a total simulation time of 1 µs. And this contact time of was used as the threshold of selection of high-water contact residue. The water pathway, comprised of the residues in contact, is illustrated in Figures 2A and 2B with the residues coloured according to their contact frequency with water in different orientations of the simulation box. Based on this analysis, water was found to traverse specific regions of the protein, namely TM 1, 2, 3, 6 and 10, with less contact observed in TM 4, 5, 7, 8 and 9 (Figure 2E). Among these residues, 10 are hydrophilic (including glutamic acid, arginine, serine, threonine, asparagine and methionine) and 5 are hydrophobic (including isoleucine, leucine and phenylalanine). Among the 11 hydrophilic residues, 8 interacted with water via their side chains, while the remaining 2 made contact through backbone atoms (Table S1). Water-contact residues identified here are not overlapping with the experimentally proven L-alanine binding pocket in GerAB that consists of G25, V101, L199, G200, T287 and Y291 ^7^. It indicates water crossing in GerAB might *not* be stabilizing the binding pocket and facilitating ligand binding as reported in GkApcT^12^, otherwise water path would go through the binding pocket. This inspired us to further verify the function of water-contact residues. Given water’s polar hydrogen-oxygen bonds and its capacity to form hydrogen bonds, we focused on verifying the functional relevance of the 8 hydrophilic residues that interact with water through their side chains. These were designated as “high-contact residues” and selected for *in silico* and *in vivo* validation.

**Figure 2.**
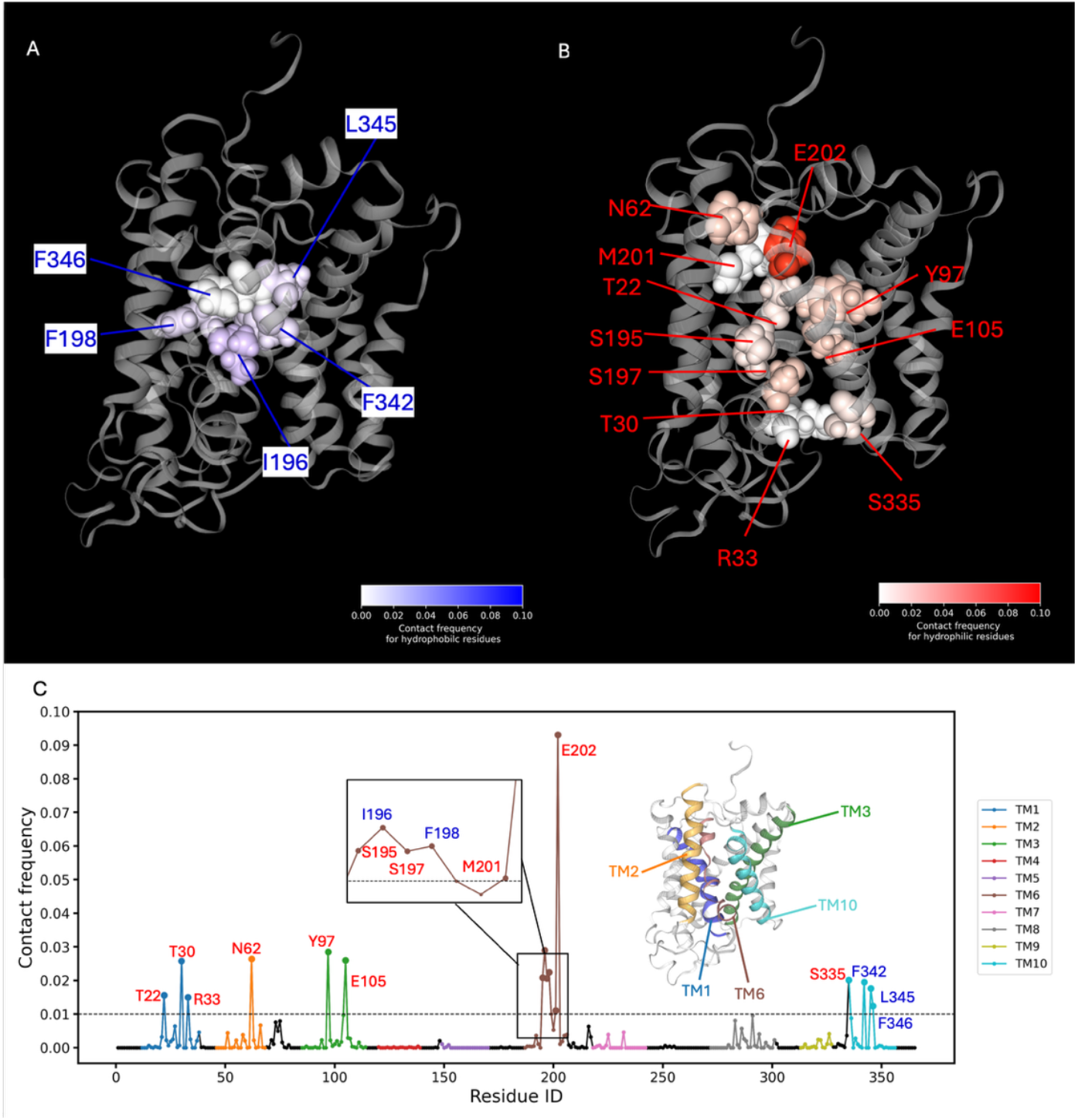
Identification of GerAB residues in contact with water. (A)-(B) High contact frequency hydrophobic/hydrophilic residues in GerAB. The protein backbone is illustrated as a grey ribbon, with high contact frequency residues highlighted in space filling according to their contact frequency *f*. Each individual residue is labelled in both subplots. (C) Contact frequency according to Residue ID and Forward frequency *f*_*F*_ are shown as a plot. The residues with a contact frequency above 0.01 in both directions are labelled. One snapshot of GerAB structure is inserted in this subplot with TM regions including high contact frequency residues represented in same colour in the plot. Hydrophilic residues are coloured in red and hydrophobic residues are coloured in blue in all subplots.

### 2.3 High contact residue alteration alters water passage profile in simulation

Based on contact frequency analysis, we first incorporated single point mutations on high-contact residues *in silico* and tracked their water passage profile in simulations. Since drastic changes in side-chain size may negatively impact the stability of GerAB as reported previously ^8,18^, target residues were mutated into residues with similar sized sidechains. This is done by computing, within each residue, which atom contacts water most frequently, represented as the percentage of total water-contact frames for that residue. The high-contact residues and their respective substitutions are illustrated in Figure 3.

**Figure 3.**
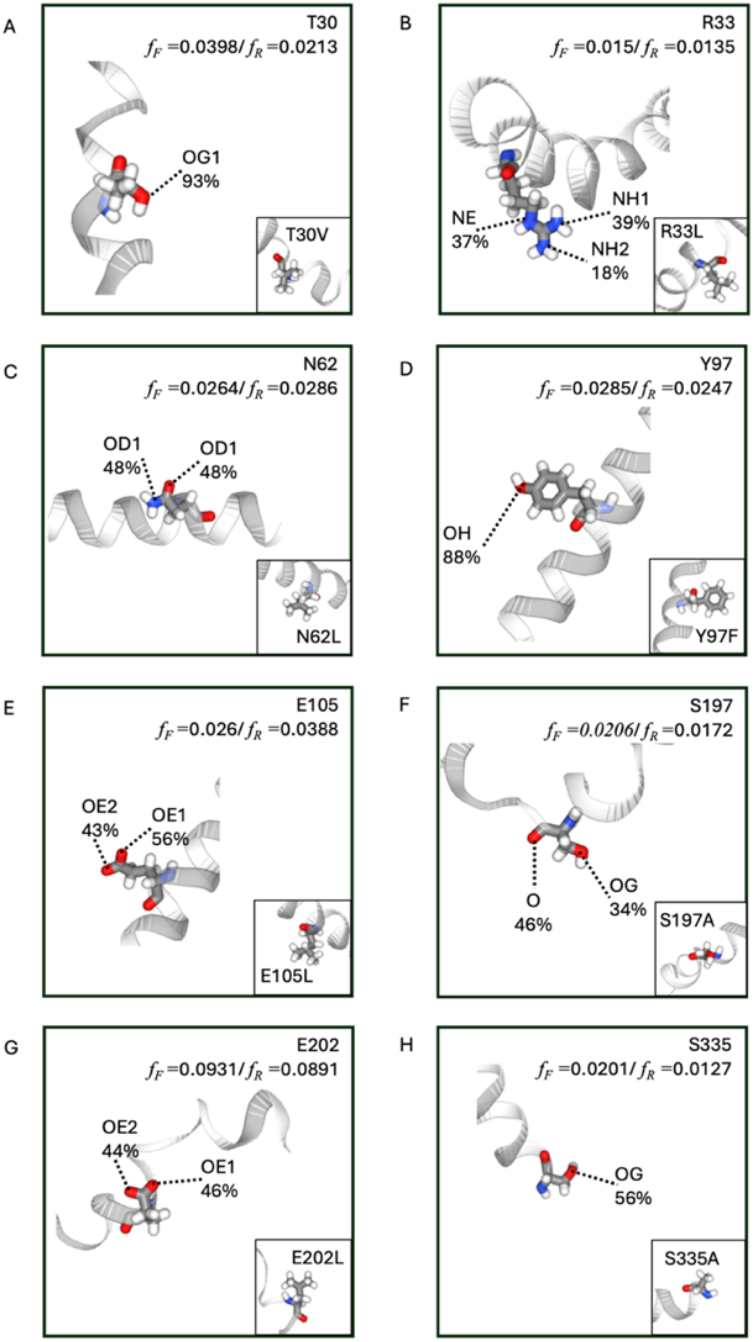
Substitution of high contact frequency hydrophilic residues. Each residue is depicted in one panel in this figure with its *f*_*F*_ and *f*_*R*_. The TM region of each residue is depicted to show the context of the protein. The protein backbone is depicted as a grey ribbon while elected residues are depicted in licorice representation, with oxygen in red, nitrogen in blue. The atom(s) with highest contribution of interactions between a residue and the passing water molecule(s) is labelled in each panel. The mutation introduced into GerAB was inserted at the bottom right corner in each panel.

We then set up MD simulations with GerAB single point mutants in the same set up as used for the wt GerAB-membrane system, see Methods for more details. Of each variant of GerAB, five replicas were run. This was followed by analysing water permeation events for each simulation run. The water passage number for each mutant simulation run are shown in Table 1 (forward direction) and Table S2 (reverse direction). Consistent with wild-type GerAB, the number of forward and reverse permeation events remained on the same order of magnitude in all systems, as the simulations were carried out under equilibrium conditions. Most of the mutants exhibited reduced water passage counts compared to the wild type simulations. Notably, some mutants, such as E202L, showed water permeation counts close to zero in both directions across all simulation replicates. This finding reinforces our analysis, as E202 was identified as the residue with the highest contact probability among all high-contact residues. On the other hand, GerAB Y97F mutant displayed an increased level of water crossing ranging from 0-164 water molecule/µs. The elevated water passage observed in the Y97F mutant may be attributed to other factors, such as altered interactions among neighbouring residues, which require further characterization.

**Table 1.**
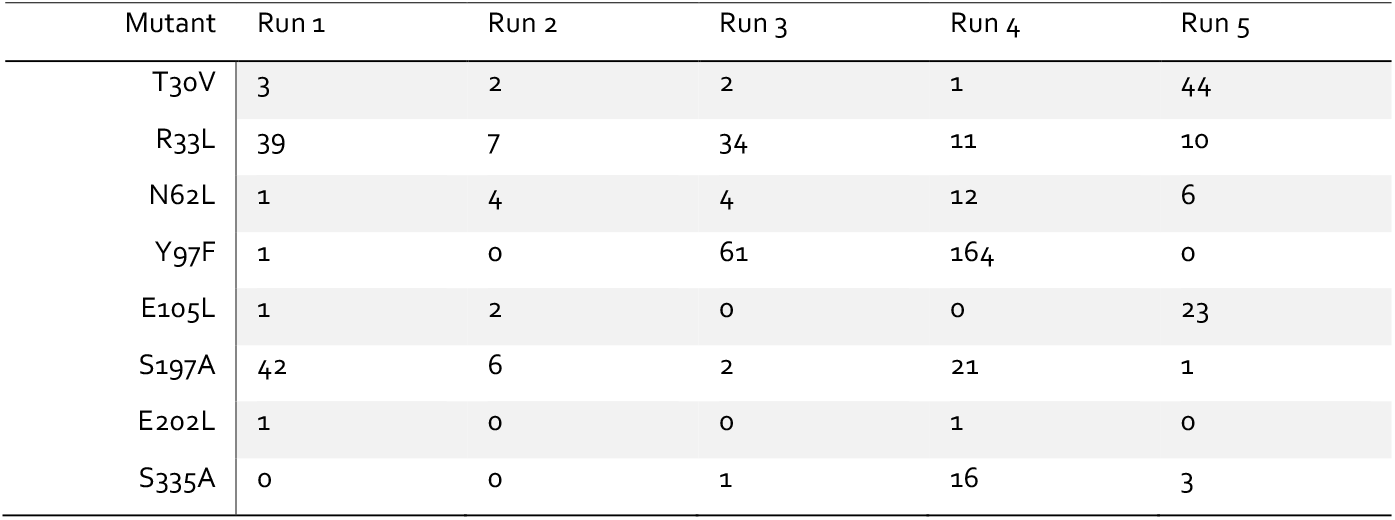
Water crossing passage number of mutant GerAB simulations in the forward direction. Each mutant shows passage number per run.

### 2.4 High contact residue alteration impairs germination via structural disruption

To verify whether these high-contact residues contribute to water permeation *in vivo*, we constructed single point mutant spores with same mutations in simulations. The effect of each mutation was verified with tracking their germination kinetics. Figure 4A shows that all 8 mutants have reduced spore viability and failed to germinate in the presence of L-alanine. To investigate whether mutant proteins were incorporated in the spore, we relied on the fact that the A and C subunits of the GerA GR depend on GerAB for stability ^18,19^. The detection of GerAA level was therefore employed as a method to detect folding of GerAB and the assembly of GerA. Immunoblot analysis showed in all the mutant spores that the levels of GerAA were similar to those of spores with Δ *gerA* (Figure 5). This means spores harbouring any of the mutants made GerAB unstable or unable to form a complex with GerAA and GerAC. To assess if making mutations on other residues with functional importance on GerAB causes structural disruption, we carried on constructing mutants in the previously reported GerAB L-alanine binding pocket, G200A and G25A^7^. Contrary to prior findings, which showed that G25A forms a stable GerA complex, we observed neither of these two binding pocket mutants showed positive GerA assembly (Figure S3). With this unexpected result, we examined the genome of the G25A mutant constructed in our lab. Our analysis showed two copies of the G25A*-gerA* operon instead of a single copy in the wt strain, indicating the gene copy number might play a determining role in GerA assembly. In summary, minor alterations to the GerAB protein on residues with high water-contact cause structural disruption of GerAB. However, changing binding pocket residue G25, which is not a high-water-contact residue, does not cause structural disruptions, according to the Western Blot in the original paper^7^. Combining that evidence, we conclude the high water-contact residues can be considered hotspots for maintaining GerAB structural integrity. With none of the high water-contact residue mutants forming a stable GerA complex, it remains still experimentally unproven whether there is water intake through GerAB.

**Figure 4.**
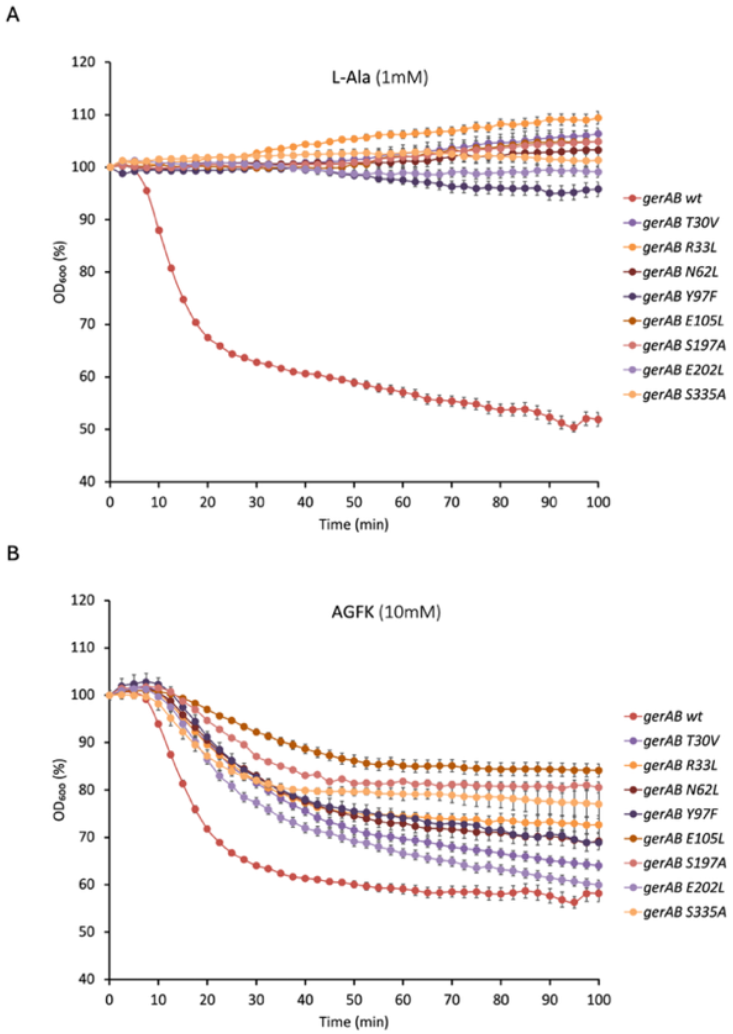
Germination assays of *B. subtilis* spores harboring wt and mutant GerAB genes with L-alanine (A) or the AGFK mixture (B). As shown in (A), all variants of mutant spores are unable to germinate with L-alanine. At the same time, all variants of mutant spores exhibited slow germination in response to the AGFK mixture, albeit to different extents (B). Error bars indicate ± SD of three technical replicates.

**Figure 6.**
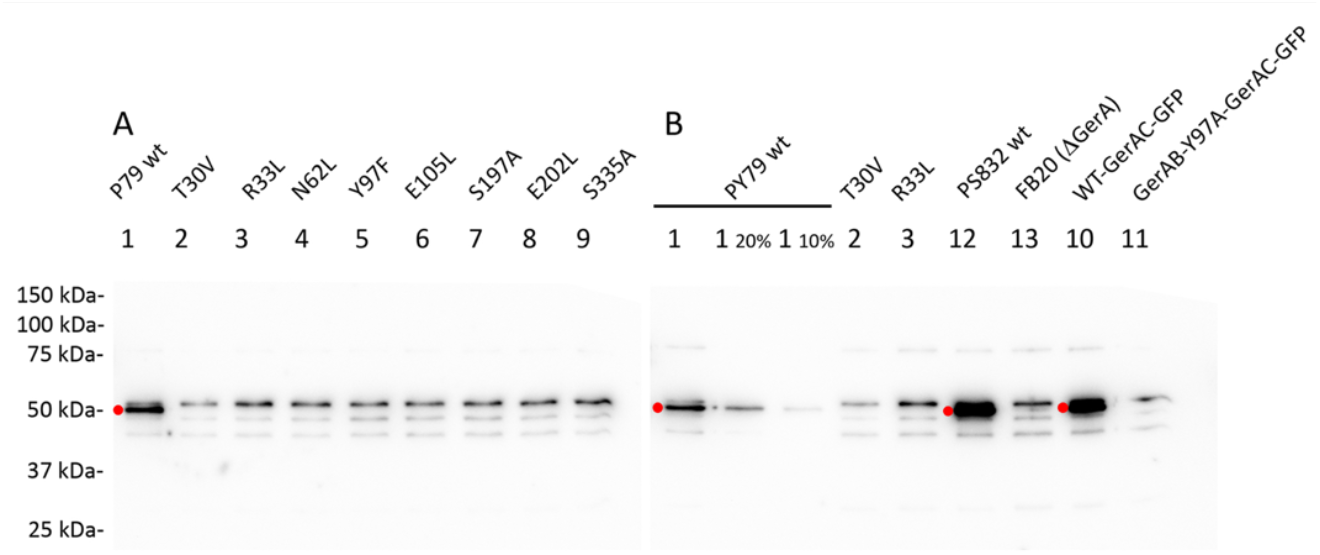
Western Blot of spores harboring different *gerAB* variant against GerAA antibody. Specific bands are around S0kDa, labelled with red dots. Noted, unspecific bands included the ones slightly above specific band. Mutant spores exhibited similar bands as spore lacking *gerA* operon, instead of with wt spore.

To further examine whether the GerAB mutations on high-contact residues impact the structure and function of other GRs in the germinosome, we performed germination assays of mutant spores against germinants that are sensed by the GerB and GerK GRs, namely the AGFK mixture. All mutants displayed reduced germination kinetics in response to AGFK, with varying degrees of impairment compared to wild-type spores (Figure 4B). Since GerB and GerK colocalize into germinosome along with GerA, this suggests the function of GerB and GerK is dependent on the structural stability and full functionality of GerA. This is also observed in a previous study ^8^, and aligns with the current understanding of germinosome function^4,5^. It is also consistent with the proposed model of subunit exchange between germinant receptors (as discussed by Prof. David Rudner at the 11th European Spore Conference, 2025), in which different GerAB mutants may assemble with GerB or GerK to varying degrees, resulting in distinct AGFK germination kinetics. However, confirming this hypothesis will require further investigation of the binding affinities between GR subunits, both within the same receptor complex and between different receptor complexes.

## 3 Discussion

With MD simulations, we identified water passage events through GerAB and further identified 8 hydrophilic residues in GerAB that have high contact frequency with water during all permeation events. Altering these residues to similar sized hydrophobic residues altered water passage *in silico*, with most of the mutants showing decreased water passage in MD simulations. The *in vivo* role of these residues was verified with mutagenesis followed by germination assay with different germinants. All 8 mutants showed severe germination defects with L-alanine and decreased germination with the AGFK mixture. With Western Blot showing no successful assembly of GerA receptors in all mutant spores, the L-alanine germination defect was caused by the absence of the GerA GR. This indicates the identified high water contact residues are also hotspots for GerAB structural integrity.

The role of water in GerAB observed in this study differs from that reported for other ApcTs, where water facilitates ligand binding^12^. A previous study identified the L-alanine binding pocket of GerAB involving G25, V101, L199 G200, T287 and Y291 ^7^, yet in our current simulation setup, the water permeation path does not pass through this predicted pocket. This discrepancy suggests water permeation in GerAB may not contribute directly to ligand binding. Given the current experiment setup of this study, we are unable to determine whether GerAB facilitate water passage to promote germination. Instead, with the constructed mutants, we found that the disruption in GerA assembly also disrupted the structure of the GerB and GerK GR which directly reflect germinosome assembly, as observed in previous study ^8^. To further prove the structural co-dependency between GRs, future studies could focus more on the affinity for the cognate A and C subunit partners in GerA, GerB and GerK.

## 4 Methodology

### 4.1 MD simulations of GerAB-membrane systems

The GerAB-membrane system was prepared according to Blinker *et al* ^10^. In short, GerAB structure was predicted by the RaptorX structural prediction tool^20^, and the structure was used subsequently to construct a protein-bilayer membrane system and introduced mutations using CHARMM-GUI ^21^, with a bilayer membrane composition mimicking the spore IM (with the lipid ratio of POPE:POPG: TMCL=1:6:3). The system was solvated by TIP3P water model and 20mM KCl was used to neutralize the system. MD simulation of the GerAB-membrane system was carried out using GROMACS engine, version 2023.3. We used the ^21,22^, and the long-distance electrostatic interactions were computed by using particle mesh Ewald (PME) algorithm^23^ with a grid spacing of 0.12 nm. The temperature was kept at 298 K with the velocity rescaling thermostat ^24^ and the pressure was kept constant around 1.0 bar using the Parrinello-Rahman barostat ^25^. To remove any steric clashes and hindrances, the systems were first energy minimized using the steepest descent algorithm ^26^, followed by equilibration carried out according to Lee et all ^21^. The production runs were 1000 ns each, using a 2 fs timestep with frames saved every 2 ps. To avoid excessively large output files, trajectories were generated in segments of 100 ns each. 10 parallel 1 µs simulations were performed under identical conditions for the wt system, and 5 parallel simulations were performed for mutant systems. Each individual simulations initiated with different random velocities. All the production runs were carried out under periodic boundary conditions (PBC). Details of the MD simulation setup are provided in Table 2.

**Table 2.**
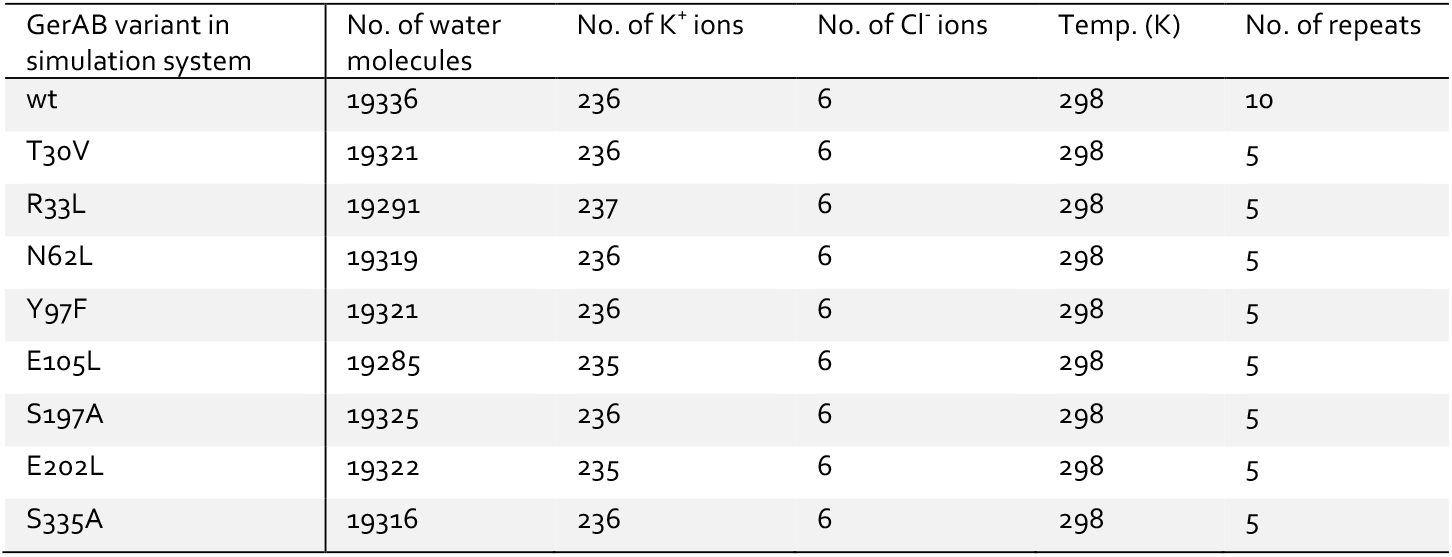
Detail of MD simulations systems.

### 4.2 Water permeation analysis

The stability of the simulation system was monitored with root-mean-square deviations (RMSD) and root-mean-square fluctuations (RMSF) of the protein carbon alpha atoms. During simulations, one water permeation event was defined as a water molecule increase in its Z coordinate in each consecutive frame after it reaches the entrance of the protein till it reaches the exit of the protein, while it remained within the cylinder of protein. If a water molecule goes through the entrance and exits in two consecutive frames (2ps), it is considered to go through periodic boundary jump and it is not considered as a valid permeation event. The entrance and the exit of the protein along the z axis was defined as Z = 2.5nm and Z = 7.5nm, respectively. The cylinder of the protein was defined between 3.5nm and 6.5nm for both x and y coordinates. Since the production runs are carried out under equilibrated conditions, water should permeate through both directions. Therefore, all the water molecules were examined in both directions and counted to give the total water passage, and all 10 runs for each system were analysed per 100ns.

### 4.3 Residue contact analysis

To determine the pathway traversed by water molecules during simulations, we computed the frequency of each residue *i* that came into proximity with permeating water, defined as *f*. To be specific, we first entailed recording the trajectory of all water molecules engaged in permeation events. For each simulation frame encompassing a water permeation event, we recorded the amino acid side chain heavy atoms within a 3.5 Å radius of the water oxygen atoms. To avoid double counting, only the closest heavy atom was counted as 1 contact. Subsequently, this count was normalized by dividing it with the total number of frames (1,285,813 in forward direction and 1,109,025 in reversed direction) encompassing all water permeation events. For each residue in contact with water, the contact frequency of each heavy atom was calculated and normalized by the total contact of that residue. The residue contact was quantified for both directions of permeation. Following the outlined procedure, all residues with water contact were visually inspected using VMD.

### 4.4 Mutagenesis

To focus on biologically relevant interactions of residues in contact while minimizing noise from transient contacts, a frequency threshold of 0.01 statistical significance was applied in residue contact analysis for both permeation direction. Resides selected for mutagenesis target are: i) hydrophilic, ii) reached 0.01 or more contact both directions, and iii) the highest contact atom is a side chain atom. For the selected residue, the atom with the highest contact was examined and the residue was altered to the structurally closest residue without the highest contact heavy atom.

All strains were derived from *Bacillus subtilis* PY79. The *gerA* operon was cloned by PCR, inserted in plasmid vector pUC19 with the Gibson Assembly Master Mix kit (New England BioLabs, NEB # E2611S). Plasmid-based mutagenesis was carried out with the QuikChange Lightning Site-Directed Mutagenesis Kit (Agilent Technologies, Cat # 210518-5). Mutant *gerAB* sequences were integrated into the original *gerAB* locus by a double crossover along with an erythromycin resistance cassette. The correct sequence of the mutant *gerAB* locus was confirmed by DNA sequencing. Mutant strains established in this study are listed in Table 3.

**Table 3.** Strains used in this study.

| Strain | Origin strain | Source |
| --- | --- | --- |
| gerAB Wild Type (wt) | *Bacillus subtilis* PY79 | Lab stock |
| gerAB T30V::erm | *Bacillus subtilis* PY79 | Constructed in this study |
| gerAB R33L::erm | *Bacillus subtilis* PY79 | Constructed in this study |
| gerAB N62L::erm | *Bacillus subtilis* PY79 | Constructed in this study |
| gerAB Y97F::erm | *Bacillus subtilis* PY79 | Constructed in this study |
| gerAB E105L::erm | *Bacillus subtilis* PY79 | Constructed in this study |
| gerAB S197A::erm | *Bacillus subtilis* PY79 | Constructed in this study |
| gerAB E202L::erm | *Bacillus subtilis* PY79 | Constructed in this study |
| gerAB S335A::erm | *Bacillus subtilis* PY79 | Constructed in this study |
| gerAB G25A::erm | *Bacillus subtilis* PY79 | gerAB S335A::erm |
| gerAB G200A::erm | *Bacillus subtilis* PY79 | Constructed in this study |
| gerAB Wild Type (wt) | *Bacillus subtilis* PS832<sup>27</sup> | Prof. Peter Setlow lab, UConn Health |
| ΔgerA | *Bacillus subtilis* PS832<sup>27</sup> | Prof. Peter Setlow lab, UConn Health |

### 4.5 Sporulation and spore purification

Bacterial strains were streaked on LB agar with the appropriate antibiotics and incubated overnight. A single colony was inoculated into liquid LB with antibiotics and grown to an OD_600_ of 1.0–2.0, after which 200 µL of culture was spread on 2xSG agar plates (Difco Nutrient Broth 16 g/L, KCl 26 mM, MgSO_4_ 2 mM, MnCl_2_ 0.1 mM, FeSO_4_ 1.08 µM, Ca(NO_3_)_2_ 1 mM, Glucose 5.5 mM, Agar 15 g/L) without antibiotics. Plates were incubated upside down at 37 °C in plastic bags for 2-5 days until sporulation, monitored by phase-contrast microscopy. Plates were then air-dried for ~ 2 days to promote cell lysis, and spores were scraped into cold Milli-Q water. Suspensions were sonicated for 1 min at full power, cooled on ice, and centrifuged at ~ 8,000 rpm for 20 min to remove debris. Spore purity was confirmed by phase-contrast microscopy.

### 4.6 Germination assay with phase contrast microscopy

Purified phase-bright spores (OD_600_ = 5 in Milli-Q water) were heat-activated at 70 °C for 30 min, then cooled on ice for 20 min. Germinant solutions were kept on ice before use. Germination was triggered by adding either 1 mM L-alanine or 10 mM AGFK (10 mM each of L-asparagine, D-glucose, D-fructose, and KCl) ^7,28^ prior to slide preparation, which followed Wen *et al* with modifications^5^. Slides and coverslips (22 mm round; 22 mm x 30 mm rectangular) were cleaned with 70% ethanol and air-dried vertically. Two rectangular coverslips were prewarmed on a 70 °C heating block, and 65 µL of 2% agar was deposited on one, then covered with the other to spread the agar evenly. After drying for ~10 min, coverslips were separated, and the agar was cut into a 1 x 1 cm patch. A 0.4 µL aliquot of spore suspension with germinant was applied to the patch, which was then transferred onto a round coverslip. The coverslip was mounted in a G-frame affixed to an air-dried slide, sealing all edges for microscopy. Time-lapse imaging was performed on a Nikon Eclipse Ti equipped with a Nikon Ti Ph3 phase contrast condenser, Nikon Plan Apo λ Oil Ph3 DM objective (100x, NA = 1.49, T = 23 °C), Lumencor Spectra Multilaser (470 nm, 555 nm) with Lambda 10-B filter blocks, NIDAQ Lumencor shutter, Ti XY- and Z-drive, and a Hamamatsu C11440-22C camera, all controlled by NIS-Elements AR 4.50.00 (Build 1117) Patch 03. The condenser was optimized before each session to yield a pixel size of 0.07 µm. For each slide, a 100 min time-lapse was recorded at 30 s intervals at 37 °C. Single-cell germination was analysed using SporeTrackerX^29^, and germination efficiency was defined as the percentage of germinated spores at the end of the time-lapse.

### 4.7 Germination assay with optical density drop

Purified phase-bright spores were normalized to an OD_600_ of 1.2 in 25 mM HEPES buffer (pH 7.4), heat-activated at 70 °C for 30 min, and cooled on ice for 20 min. Aliquots (100 µL) of heat-activated spores were dispensed into a 96-well plate, and germinants were added to final concentrations of 1 mM L-alanine or 10 mM AGFK. OD_600_ was recorded every 2.5 min for 100 min at 37 °C with continuous agitation between readings. OD reduction was calculated relative to the initial value. Each assay was performed in triplicate, and mean OD drops are reported.

### 4.8 Germination assay with DPA release

CaDPA release was quantified using a fluorescence plate reader as described by Yi and Setlow^30^. Spores (OD_600_ = 0.5) were germinated with 1 mM L-alanine in 25 mM K-HEPES buffer (pH 7.4) at 45 °C without prior heat activation. At designated time points, 190 µL of the suspension was mixed with 10 µL of 1 mM TbCl3, and relative fluorescence units (RFU) were recorded.

### 4.9 SDS-PAGE and immunoblotting

For western blotting, spores were decoated and lysed as follows. 50 OD units of spores were pelleted by centrifugation, resuspended in 1 mL TUDSE buffer (8 M urea, 50 mM Tris-HCl pH 8.0, 1% SDS, 50 mM DTT, 10 mM EDTA), and incubated at 37 °C for 45 min. After centrifugation (3 min, max rpm, room temperature), the pellet was resuspended in 1 mL TUDS buffer (8 M urea, 50 mM Tris-HCl pH 8.0, 1% SDS) and incubated again for 45 min at 37 °C. The spores were washed six times by centrifugation and resuspended in 1 mL TEN buffer (10 mM Tris-HCl pH 8.0, 10 mM EDTA, 150 mM NaCl). Decoated spores were stored in water if not processed immediately. For lysis, 50 OD units of decoated spores were treated with 1 mg lysozyme in 0.5 mL TEP buffer (50 mM Tris-HCl pH 7.4, 5 mM EDTA) containing 1 mM PMSF, 1 µg RNase, 1 µg DNase I, and 20 µg MgCl_2_ at 37 °C for 6-8 min, followed by 20 min on ice. Glass disruptor beads (0.10—0.18 mm, 100 mg) were added, and spores were sonicated in three 10 s bursts (medium power, microprobe) with 30 s cooling on ice between bursts. After settling for 15 s, 100 µL of supernatant was mixed with 100 µL 2x Laemmli buffer (5% 2-mercaptoethanol, 1 mM MgCl_2_) and boiled at 90-95 °C for 3 min to obtain the total lysate. Twenty micrograms of total protein per lane were separated on a 10% SDS-PAGE gel (BIO-RAD #4561034) at 60 V for 45 min, then 110 V for 45 min, and transferred to a 0.22 µm PVDF membrane. The membrane was blocked with 2.5% low-fat milk in TBST (20 mM Tris, 150 mM NaCl, 0.1% Tween-20) for 30 min, incubated overnight with anti-GerAA antibody (1:3000) (20), washed three times with TBST, and probed with goat anti-rabbit HRP antibody (1:2500; BIO-RAD #1706515) for 1 h before visualization.

## Supporting information

https://doi.org/10.6084/m9.figshare.31042450

## Notes

### Competing Interest Statement

The authors have declared no competing interest.

### Funding

- China Scholarship Council: 202109120009

