## Supplementary material for "GerAB Residues Predicted to Interact with Water Based on MD Simulations Mediate Germinosome Stability in *Bacillus subtilis* spores": https://doi.org/10.6084/m9.figshare.31042450

### Supplementary Information

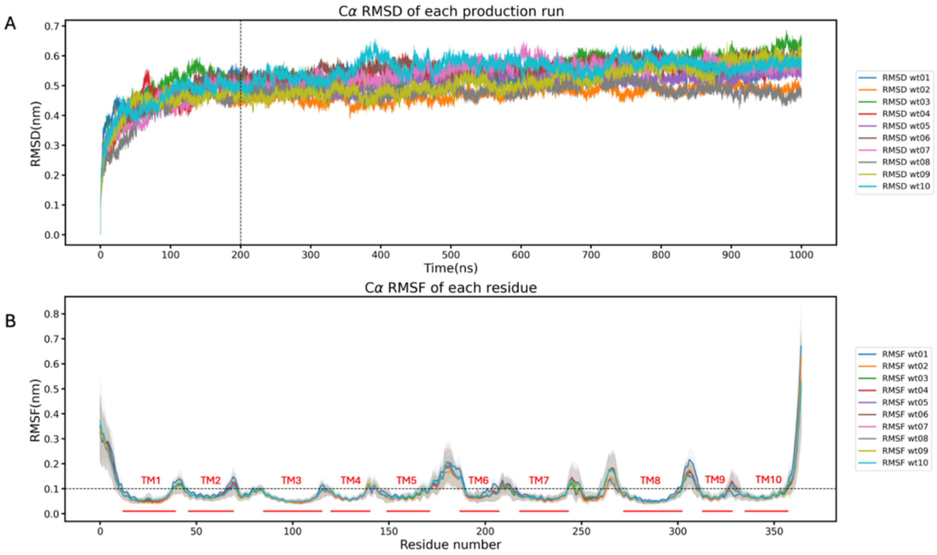


Figure S1. Protein structural stability during ten parallel MD simulations production runs. Protein Ca RMSD (A) increased from 0 to 200ns simulations and stabilized afterwards at around 0.6nm in all 10 simulation runs. Protein Ca RMSF (B) showed the RMSF of each residue calculated per 100ns and averaged over each production run. Error margin showed std of RMSF of each residue. Ten production runs showed consensus over stable regions (helix) and high-fluctuation regions (loops). All TM regions were marked as red bars in the figure. Both analyses showed protein remains structurally stable during all simulation runs.

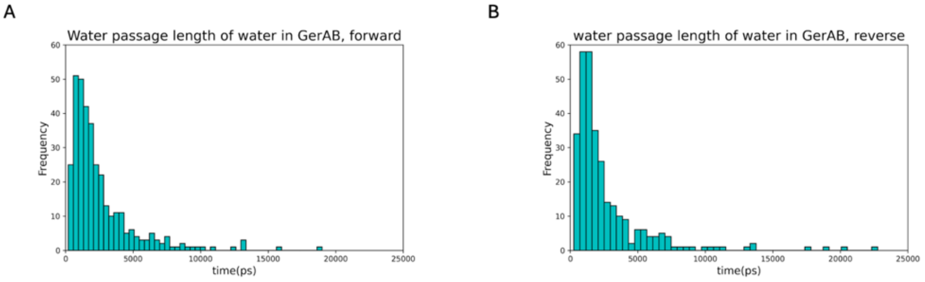


Figure S2. Distribution of the length of the water passage events. Most of the water passages are below 5ns.

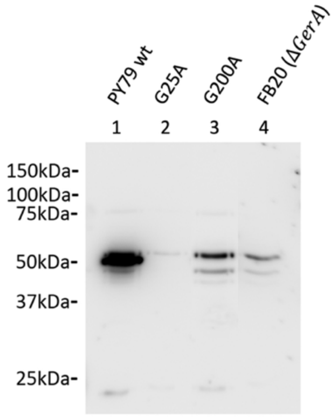


Figure S3. Western Blot of spores harboring different gerAB variant against GerAA antibody. Both G25A and G200A spores exhibited similar bands as spore lacking *gerA* operon. Notably G25A mutants was reported to exhibit positive GerAA cross reaction material. Notably the G25A mutant strain tested here includes two copies of *gerA* operon. This figure shows partial Western Blot from one experiment; irrelevant samples were cropped for clarity.

| *Residue* | *Mutation* | $f_F/f_R$ | *atomF* | *atomR* |
|---|---|---|---|---|
| *E202* | **E202L** | 0.0931/0.0891 | OE1 44%; OE2 46% | OE1 49%; OE2 40% |
| *I196* | N/A | 0.0290/0.0229 | O 95% | O 96% |
| *Y97* | **Y97F** | 0.0285/0.0247 | OH 88% | OH 88% |
| *N62* | **N62L** | 0.0264/0.0286 | ND2 49%; OD1 48% | ND2 36%; OD1 60% |
| *E105* | **E105L** | 0.0260/0.0388 | OE1 56%; OE2 43% | OE1 53%; OE2 47% |
| *T30* | **T30V** | 0.0258/0.0213 | OG1 93% | OG1 94% |
| *F198* | N/A | 0.0224/0.0166 | O 97% | O 98% |
| *S195* | N/A | 0.0208/0.0113 | O 92% | O 88% |
| *S197* | **S197A** | 0.0206/0.0172 | O 65%; OG 34% | O 80%; OG 19% |
| *S335* | **S335A** | 0.0201/0.0127 | OG 56%; O36% | OG 55%; O 40% |
| *F342* | N/A | 0.0195/0.0184 | O 62%; CE2 13% | O 57%; CE2 17% |
| *L345* | N/A | 0.0176/0.0130 | O 88% | O 83% |
| *T22* | N/A | 0.0156/0.0250 | O 83%; OG1 17% | O 58%; OG1 41% |
| *R33* | **R33L** | 0.0150/0.0135 | NH2 39%; NE 37%; NH1 18% | NH2 35%; NE 41%; NH1 21% |

Table S1. High contact frequency ($f$) residues identified in this study. Only both $f_F$ and $f_R$ higher than 0.01 were listed in this table, as shown in Figure 2 in main text. AtomF and AtomR listed the heavy atoms on each residue side chain that interact with the water oxygen on direction forward and reverse, respectively. O is backbone oxygen while OG, OE, OD, OH are sidechain oxygen. NH, NE, ND and CE are sidechain heavy atoms. The atom label follows that of a standard PDB file.

| Mutant | Run 1 | Run 2 | Run 3 | Run 4 | Run 5 |
|---|---|---|---|---|---|
| T30V | 0 | 3 | 4 | 1 | 44 |
| R33L | 47 | 7 | 30 | 8 | 5 |
| N62L | 1 | 0 | 1 | 7 | 2 |
| Y97F | 1 | 4 | 54 | 151 | 2 |
| E105L | 1 | 1 | 0 | 0 | 24 |
| S197A | 56 | 0 | 3 | 30 | 4 |
| E202L | 2 | 1 | 1 | 2 | 0 |
| S335A | 1 | 0 | 0 | 18 | 1 |

Table S2. Water crossing passage number of mutant GerAB simulations in the reverse direction. Each mutant shows passage number per run.
